# The amino acid residue in position 163 of canine PrP^C^ is critical to the exceptional resistance of dogs to prion infections: evidence from transgenic mouse models

**DOI:** 10.1101/800698

**Authors:** Enric Vidal, Natalia Fernández-Borges, Hasier Eraña, Beatriz Parra, Belén Pintado, Manuel A Sánchez-Martín, Jorge M Charco, Montserrat Ordoñez, Miguel A Pérez-Castro, Martí Pumarola, Candace K. Mathiason, Tomás Mayoral, Joaquín Castilla

**Author notes:** These authors contributed equally to this work (Enric Vidal, Natalia Fernández-Borges). To whom correspondence should be addressed. Joaquín Castilla, CIC bioGUNE, Parque tecnológico de Bizkaia, Derio 48160, Bizkaia, Spain, **E-mail:**.

## Abstract

Unlike other species, such as cattle, cats or humans, prion disease has never been described in dogs, even though they were similarly exposed to the bovine spongiform encephalopathy (BSE) agent. This resistance prompted a thorough analysis of the canine *PRNP* gene and the presence of a negatively charged amino acid residue in position 163 was readily identified as potentially fundamental as it differed from all known susceptible species. Furthermore, recent results from our group demonstrated that mouse *PRNP* with the dog substitution N158D (mouse equivalent to position 163) rendered mice resistant to prion infection. In the present study, a transgenic mouse model was generated expressing dog prion protein (with glutamic acid at position 163) and challenged intracerebrally with a panel of prion isolates (including cattle BSE, sheep scrapie, atypical sheep scrapie, atypical BSE-L, sheep-BSE and chronic wasting disease, among others) none of which could infect them. The brains of these mice were subjected to *in vitro* prion amplification and failed to find even minimal amounts of misfolded prions providing definitive experimental evidence that dogs are resistant to prion disease. Subsequently, a second transgenic model was generated in which aspartic acid in position 163 was substituted for asparagine (the most common amino acid in this position in prion susceptible species) and this mutation resulted in susceptibility to BSE-derived isolates.

These findings strongly support the hypothesis that the amino acid residue at position 163 of canine PrP^C^ is a major determinant of the exceptional resistance of the canidae family to prion infection and establish this as a promising therapeutic target for prion diseases.

**AUTHOR SUMMARY:** Cats, cattle, people and dogs were all exposed to mad cow disease but, unlike the other three, dogs never succumbed to the disease. We generated a mouse model expressing canine prion protein (instead of mouse prion protein) to provide experimental evidence that dogs are resistant to prion infection by challenging the mice with a panel of prion isolates. None of the prions could infect our transgenic mice that expressed dog prion protein. When the prion protein amino acid sequence of dogs was compared to that of other susceptible species, one amino acid in a specific position was found to be different to all the prion-susceptible animals. To determine if this amino acid was the one responsible for dogs’ resistance to prions, a second mouse model was generated with the canine prion protein but the critical amino acid was substituted for the one susceptible species have. When this model was challenged with the same panel of prions it could be infected with at least one of them. These results demonstrate the relevance of this amino acid position in determining susceptibility or resistance to prions, and this information can be used to design preventative treatments for prion diseases.

## INTRODUCTION

Prion diseases are a group of invariably fatal neurodegenerative disorders for which no effective treatment or prophylaxis exist currently. Many mammalian species are susceptible and all share a common pathogenesis: the misfolding of the host-encoded cellular prion protein (PrP^C^) into a pathological conformer (PrP^res^) that accumulates in the brain leading to neurodegeneration and death[1–3]. Research efforts have been directed primarily at human prionopathies and those of domestic animals of commercial interest. However, other species have been of interest either as a disease model or due to their lack of susceptibility to infection. The study of species with significantly different prion susceptibilities is key to understanding the biological mechanisms underlying these diseases.

The PrP^C^ misfolding event can be sporadic (putatively spontaneous), caused by mutations in the *PRNP* gene or triggered by externally acquired infectious prions. Currently, the “mad cow disease” epizootic is under control but other animal prion diseases, such as scrapie in small ruminants or chronic wasting disease (CWD) in cervids, are endemic in many countries and the recent spread of CWD to the European continent is of great concern[4,5]. Interspecies transmission of prions is a well-established phenomenon and bovine spongiform encephalopathy (BSE) is one of the best examples. Exposure of various species to feedstuff contaminated with BSE prions caused several diseases including variant Creutzfeldt-Jakob disease (vCJD) in humans[6], feline spongiform encephalopathy (FSE) in domestic cats[7] and BSE in goats[8], to name a few. Therefore, the risk that this might occur with other prion diseases and cohabiting host species must not be neglected especially considering spontaneous cases of prion disease have been reported worldwide in humans[9], cattle[10] and small ruminants[11], and may exist in other species that have not been as extensively examined for prion diseases.

In some species, despite having been exposed to prions, no field cases of prion disease have ever been diagnosed. This may be for many reasons including; a low number of individuals examined, a short lifespan resulting in death from other causes before any prion disease can develop, culling at an early age or other circumstances might explain why species that have been proven susceptible to prion disease experimentally have never had naturally occurring cases reported. These include pigs[12–15], rabbits[16], mice, non-human primates[17–19], ferrets[20] and even horses where a transgenic mouse model with equine PrP was used[21].

Dogs are not included in this list for two reasons; (1) experimentation on dogs using prions is very limited (for various reasons, including ethical constraints) and (2) no transgenic model has been generated. To date, no evidence exist that dogs can be infected naturally with prions, only theoretically using an *in vitro* assays that have, under extreme and specific conditions, succeeded in misfolding dog PrP^C^[22].

To complicate things further, prions can misfold into well differentiated conformations with specific pathobiological features, the prion strain phenomenon[23]. Specific species are susceptible to a particular prion strain depending on the compatibility between the host PrP amino acidic sequence and the strain conformation of the infecting prion: for example, cats, despite having a PrP amino acidic sequence very similar to dogs, can be readily infected with BSE[24] and CJD[25] but only with great difficulty using CWD[26] (incubation period over three years). So when assessing susceptibility of any species to prions not only is the host’s PrP^C^ sequence important but also the strain of prion. The theoretical susceptibility can be predicted by examining the misfolding capability of the chosen species’ PrP^C^ *in vitro* by protein misfolding cyclic amplification (PMCA)[27]. Rabbits, a species with no reported field cases of TSE despite being sympatric with several prion susceptible ruminant species, were shown to have PrP^C^ that was readily misfolded by BSE *in vitro*[22], and susceptibility to this prion strain was further corroborated by bioassay in transgenic mice with rabbit *PRNP*[28] and experimentally *in vivo*[16]. However, the scenario in horses, another putatively prion-resistant species, is somewhat different as horse PrP^C^ can be misfolded, either *in vitro* by PMCA or by means of bioassay in mice expressing equine PrP^C^ (TgEq) (albeit with low efficiency), but the resultant horse-adapted prions are unable to propagate disease in TgEq mice, even though their ability to infect the original species remains unaltered. This is interpreted as a non-adaptive prion amplification (NAPA) phenomenon[21].

We have demonstrated that wild type dog PrP^C^ (with an aspartic acid in position 163) could be misfolded by BSE prions *in vitro* by PMCA and the resultant prions were infectious in TgBov mice[22,29] but there is still no evidence *in vivo* of PrP^res^ propagation in dogs. This resistance to prion disease makes canids, particularly the domestic dog (*Canis lupus familiaris*), an interesting species to study as, although having been exposed to BSE contaminated feed like cats, no definitive field case has ever been published despite a few unconfirmed reports[30,31].

Sequence alignment studies of the *PRNP* gene identified the presence of either glutamic (E) or aspartic (D) acids (both negatively charged amino acids) in position 163 in dogs PrP^c^ when compared to cats and these might be responsible for the differing resistance of the two species with respect to susceptibility to BSE[24] and CWD[26]. Furthermore, mouse *PRNP* with substitutions equivalent to the canine amino acid residues proved to be resistant to conversion to PrP^res^ both *in vitro*, by means of recombinant PrP-based PMCA, and *in vivo* in two different transgenic mouse models with asparagine (N) to aspartic acid substitution at position 158 (N158D)[32]. Additionally mouse *PRNP* with this canid substitution provided a protective dominant negative effect by inhibiting PrP^C^ conversion in transgenic chimeras co-expressing wild type (WT) mouse *PRNP*[33]. The same substitution introduced into a bank vole PrP^C^ transgenic mouse model significantly delayed prion propagation in this highly prion susceptible model[34]. All members of the *Canidae* family share a virtually identical *PRNP* sequence with only a few polymorphic variants present. Amongst those, the presence of aspartic acid (D) and glutamic acid (E) in position 163 stands out as it is almost exclusive to this family[35] which may be a possible evolutionary advantage as their diet is frequently based on ruminant meat [36],[37].

In the present study, a transgenic mouse line has been generated bearing wild type E163 dog *PRNP* and challenged with a variety of prion isolates. To prove that the presence of a negatively charged amino acid at position 163 in canine PrP^C^ is critical in determining resistance to prion disease, one additional transgenic mouse line was generated expressing dog *PRNP* but with asparagine 163 (D163N) as this residue at this position is present in most of the prion susceptible species. This model was then exposed to the same panel of isolates.

In this study, we confirm for the first time that dog PrP^C^ is unable to propagate any of the prion isolates we challenged them with definitively showing that canids are highly resistant to prion infection and that the resistance mechanism is encoded by the amino acid present at position 163 (D/E in canines vs. N in the rest of prion susceptible species).

## RESULTS

### Generation of *TgDog E163*: a model to evaluate canine susceptibility to prion infection

Once the effects of D/E at position 163 on mouse *PRNP* had been established[32,33] the next logical experiment was to test the prion susceptibility in an *in vivo* model bearing wild type canine *PRNP*. Considering obvious ethical and budgetary restrictions of using dogs as model, a transgenic mouse approach was pursued. Based on our previous experience, new mouse lines were generated by pronuclear injection of a construct consisting of the mouse PrP promoter and the E163 dog *PRNP* sequence. Six founders were obtained that transmitted the transgene to their progeny. After backcrossing to a line that did not express endogenous PrP (STOCK-Prnptm2Edin), expression levels of the transgene were analyzed by Western blot and one line was excluded because it expressed 10 times the levels of the endogenous gene and this could cause an undesired PrP^C^ over-expression associated phenotype[38,39]. Four lines expressed less than 2 times the endogenous gene levels and were also excluded (additionally, two of those did not breed efficiently). Finally, only hemizygous line *TgDog E163* (line 014) reproduced well and showed a consistent expression pattern of 2x compared to the endogenous dog prion protein level with an unaltered glycoform ratio upon Western blotting (Supplementary information, Fig. S1). Moreover, the immunohistochemical labelling pattern of PrP^C^ was comparable to that of a wild type mouse (Supplementary information, Fig. S1B). This line was selected for further studies.

### *TgDog* PrP^C^ *in vitro* and *in vivo* misfolding studies; none of the prion isolates resulted in misfoldin

#### *TgDog E163 in vitro* studies

An attempt was made to misfold dog PrP^C^ by PMCA using *TgDog* brain homogenates as substrates and using different prion strains as seeds. Ten rounds of serial PMCA were performed using 4 replicates for each seed including: cattle classical BSE (BSE-C), BSE-L, sheep-BSE, sheep scrapie, atypical scrapie, mule deer CWD, experimental feline CWD and BSE dog(D163)-PrP^res^ (inoculum obtained *in vitro* by PMCA using dog (D163) brain homogenate as a substrate and cattle BSE as a seed)[22]. None of the isolates tested was able to misfold *TgDog E163* PrP^C^ (Supplementary information, Fig. S2).

#### *TgDog E163* bioassay

Even though *in vitro* results usually correlate well with bioassay, ultimately, infectivity can only be demonstrated by *in vivo* inoculation. The isolates used for inoculation were the ones described in the *in vitro* section above and negative control inocula were also included consisting of normal brain homogenates (NBH) from cattle, dog and sheep.

None of the animals showed neurological clinical signs compatible with a TSE. Table 1 shows the number of animals inoculated for each isolate and a range of survival times post inoculation. Prion disease was ruled out in all the animals studied by means of Western blotting for detection of PrP^res^, histopathology and PrP^res^ immunohistochemistry.

**Table 1:**
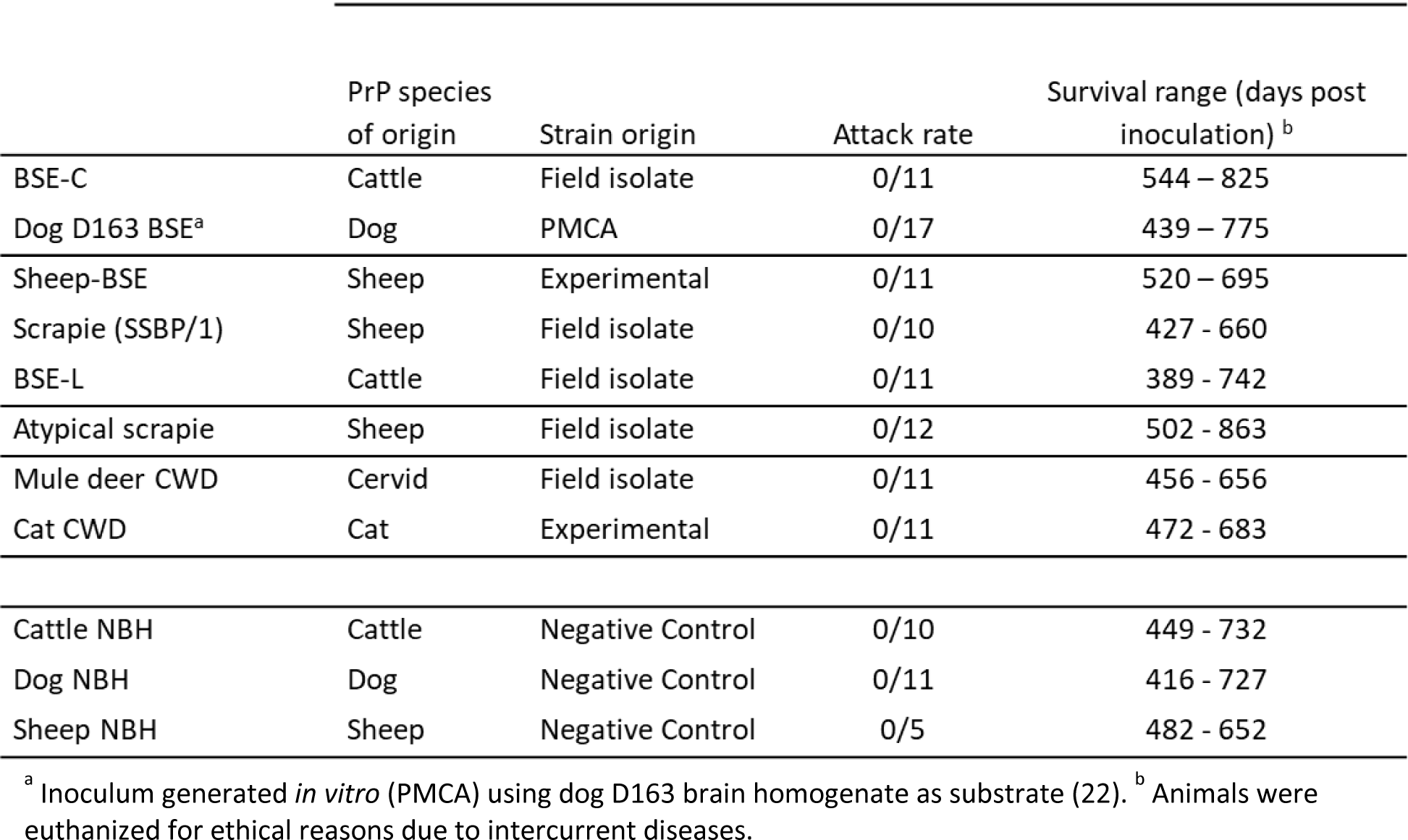
Attack rates and survival times of the inoculated *TgDog E163* mice

### PMCA propagation in brains of inoculated *TgDog E163*; unsuccessful propagation of PrP^res^ confirmed

Even though no clinical signs nor PrP^res^ deposits were detected in any of the inoculated *TgDog E163* mice by standard PrP^res^ detection methods [Western blot (WB) and immunohistochemistry (IHC)], PMCA was performed using perfused *TgDog E163* brain homogenates as a substrate to determine if even a minute amount of PrP^res^ was present that could indicate otherwise undetected *in vivo* prion protein misfolding. Pools of brains of *TgDog* mice inoculated for the bioassay study were prepared and used as seeds in the PMCA experiments. Six serial rounds of PMCA were performed to ensure the detection of minimal amounts of PrP^res^ and thereby rule out a putative propagation on a second *in vivo* passage. None of the pools showed detectable PrP^res^ after the six *in vitro* propagation rounds (Supplementary information, Fig. S3).

### Generation of *TgDog D163N* mice: a model to determine the protective effect of aspartic acid at position 163 of the dog PrP^C^

Since *TgDog E163* mice were unable to propagate prions, as shown in previous studies with mouse and bank vole PrP^C^ [32–34], the amino acid that conferred apparent resistance, aspartic acid, was removed and substituted by asparagine at positon 163 to determine if susceptibility to prions was recovered. New mouse lines were generated by pronuclear injection of a construct consisting of the mouse PrP promoter and the dog PrP sequence with the D163N substitution. From a total of 5 positive animals, 4 animal founders transmitted the transgene to their progeny. After backcrossing to a line that did not express endogenous PrP (STOCK-Prnptm2Edin), expression levels of the transgene were analyzed by Western blot. One line expressed less than 1x the wild type dog PrP^C^ levels and was discarded, another line was discarded because it expressed 5x the dog PrP^C^ levels and there was a risk of an overexpression phenotype. Of the two remaining lines, *TgDog D163N* (Line 483), expressing 2x the levels of dog PrP^C^ and with conserved glycoform ratio upon Western blotting, was chosen since this was the overexpression level obtained with the previous model (*TgDog E163*) (Supplementary information, Fig. S4A). Furthermore, immunohistochemical labelling of PrP^C^ was comparable to that of a wild type mouse (Supplementary information, Fig. S4B).

### *TgDog D163N* mice are susceptible to classical BSE and sheep-BSE *in vitro* and *in vivo*

*In vitro* studies: PMCA was performed using *TgDog D163N* mouse brain homogenates as a substrate. The same isolates as in previous sections were used as seeds and were subjected to 10 serial PMCA rounds with 4 replicates each. In contrast to what happened with *TgDog E163* brain homogenates, classical BSE and sheep-BSE were successfully propagated in this substrate (Supplementary information, Fig. S5). This result suggests that the amino acid residue substitution D163N was responsible for the recovered susceptibility to PrP^C^ misfolding.

*TgDog D163N bioassay:* Bioassays were conducted to ascertain if *TgDog D163N* mice were susceptible to prion infection in agreement with the *in vitro* results. The same panel of isolates mentioned above was used (Table 2). None of the inoculated mice developed TSE-associated clinical signs, although very mild clinical signs might have been masked by age-related changes. However, upon euthanasia 6/11 mice inoculated with sheep-BSE showed evidences of infection as confirmed by Western blotting and/or immunohistochemistry (Fig. 1). These data support the *in vitro* results that mutated dog PrP^C^ (D163N) is more susceptible to misfolding than wild type dog PrP^C^.

**Table 2:**
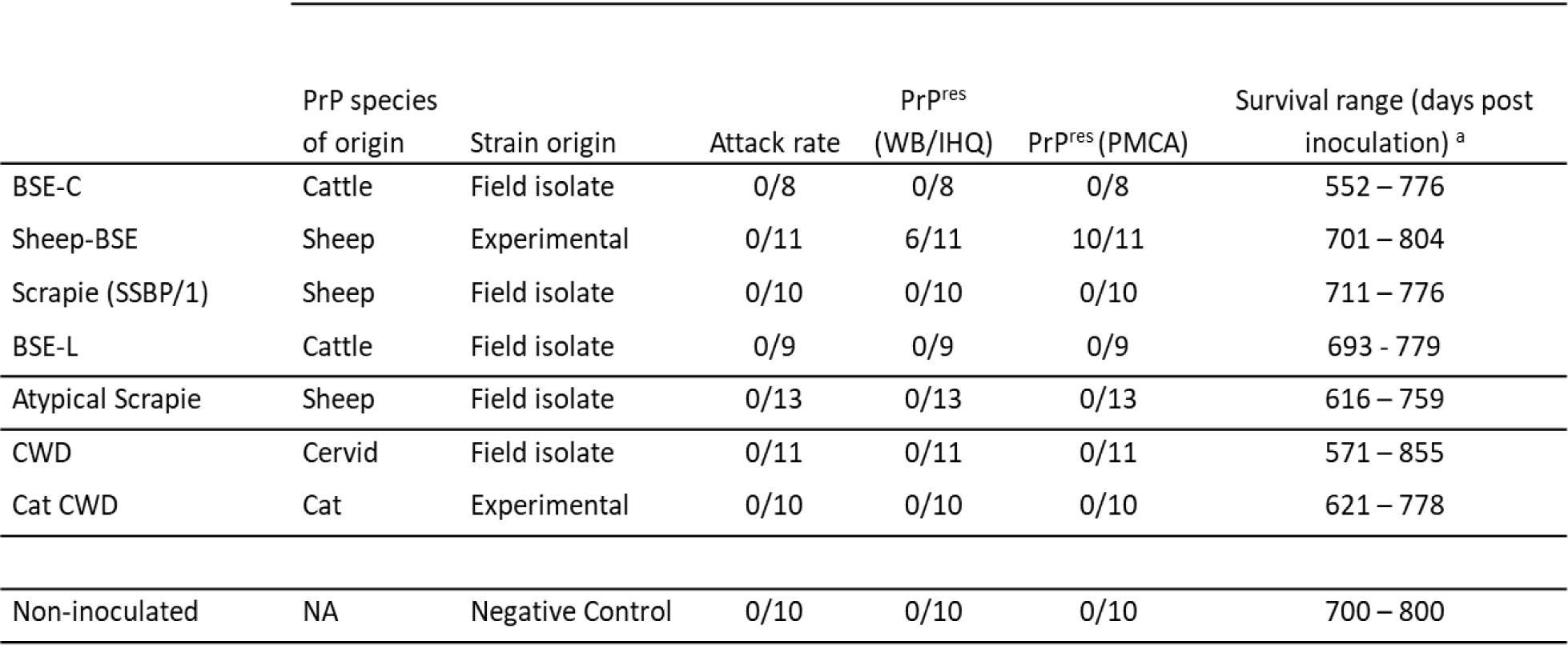
Attack rates and survival times of the inoculated *TgDog E163N* mice

**Fig. 1:**
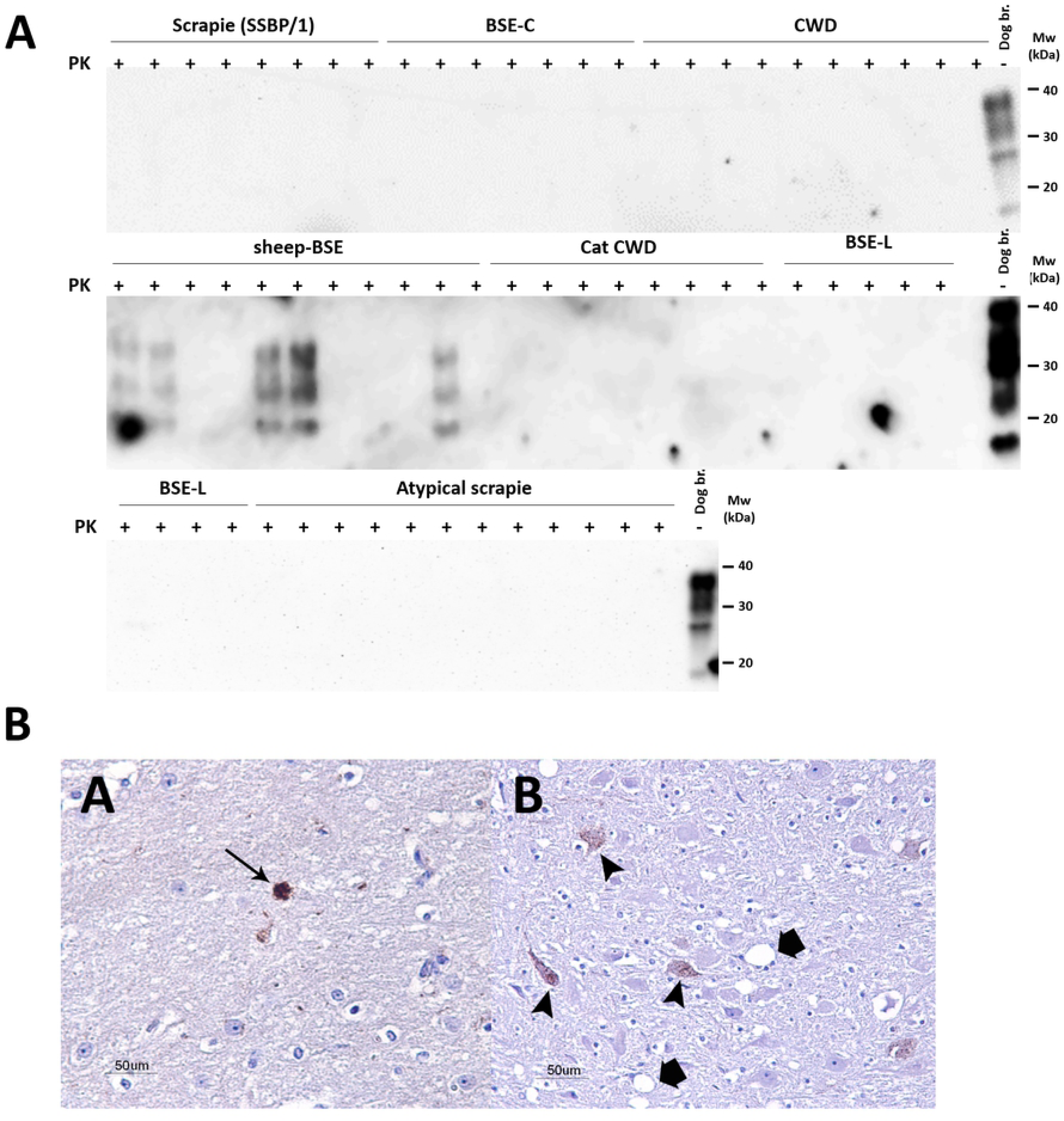
A: Molecular and pathological study of *TgDog D163N* mice inoculated with different prion isolates. **A.** Biochemical analysis of Protease-K (PK) resistant PrP (PrP^res^) in brain homogenates from *TgDog D163N* inoculated with scrapie (SSBP/1), BSE-C, mule deer CWD, sheep-BSE, cat CWD, BSE-L and atypical scrapie. The *TgDog D163N* brain homogenates were digested with 200 µg/ml of PK and analyzed by Western blot with monoclonal antibody D18 (1:5,000). Only 5 out of 10 animals inoculated with sheep-BSE showed the classical three-banded pattern after PK digestion. Dog br.: Undigested *TgDog D163N* brain homogenate. Mw: Molecular weight. **B:** Histological evidence of prion disease in brains of *TgDog D163N* mice inoculated with sheep-BSE. Panel A: Plaque like PrP^res^ deposits (brown pigment) in the thalamus (arrow). Panel B: Intraneuronal PrP^res^ deposits (brown pigment) in the thalamus (arrowheads). Note moderate spongiform change (thick arrows). PrP^res^ immunohistochemistry (mAb 6H4, 1:100, Prionics AG.).

### *In vitro* amplification of potentially undetected PrP^res^ in *TgDog D163N* mouse brains

In order to rule out that any of the other isolates inoculated in *TgDog D163N* had propagated in minute amounts undetectable by standard PrP^res^ detection techniques but could be transmitted on a second passage, PMCA was performed using *TgDog D163N* mouse brain homogenates as substrate and pooled brains from each group of inoculated mice as seeds. Each pool was subjected to 6 rounds of serial PMCA providing PrP^res^ detection sensitivity comparable to, if not greater than, a 2^nd^ passage *in vivo*[40]. In this case the sheep-BSE inoculated mice brain pool served as a positive control.

With the exception of sheep-BSE inoculated mice, no PrP^res^ propagated in any of the remaining brain pools (Fig. 2A and Supplementary information, Fig. S6). Serial PMCA was then repeated individually with the brains of mice inoculated with sheep-BSE in which no PrP^res^ had been detected by WB or IHC. Of these animals, 10 out of 11 had PrP^res^ present after *in vitro* amplification confirming the effectiveness of the PMCA procedure to reveal subclinical prion infections on 1^st^ passage bioassay (Fig. 2B). Homogenates from the mouse brains inoculated with cattle BSE were also tested individually by serial PMCA and all of them failed to propagate PrP^res^ (Fig. 2B) confirming the high specificity of the method.

**Fig. 2:**
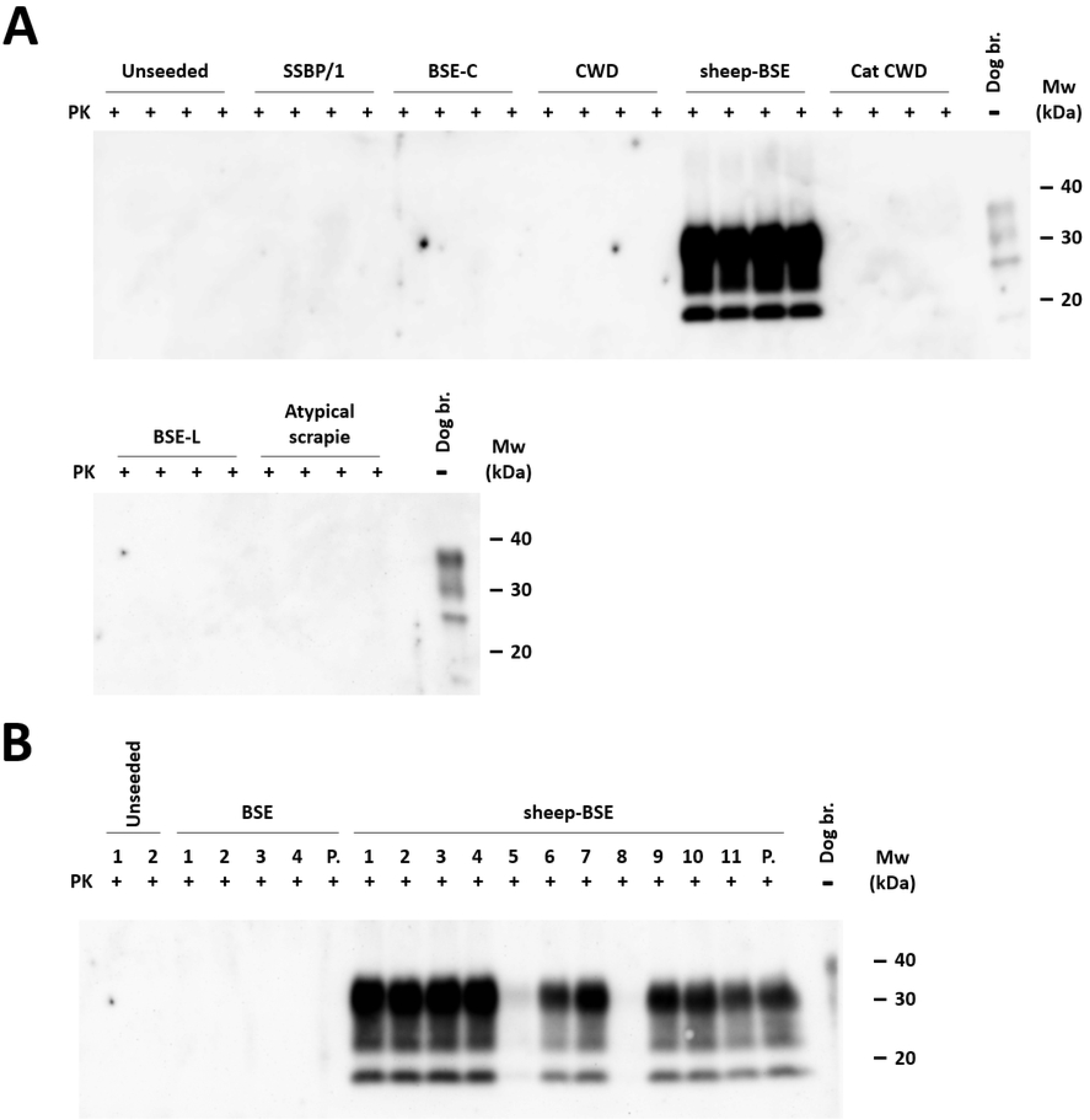
*In vitro* propagation of brain samples from *TgDog D163N* mice inoculated with different prion isolates using brain-based PMCA. **A.** 3 rounds of serial PMCA using *TgDog D163N* brain homogenates as substrate. Pools of brain samples from *TgDog D163N* inoculated with scrapie (SSBP/1), BSE-C, mule deer CWD, sheep-BSE, cat CWD, BSE-L and atypical scrapie were used as seeds in replicates of four through 3 rounds of serial PMCA. Only brain homogenates from sheep-BSE-inoculated mice propagated and this showed a classical three-banded glycosylation pattern. **B.** Individual brains of *TgDog D163N* mice inoculated with BSE-C and sheep-BSE were subjected to 3 rounds of serial PMCA using *TgDog D163N* brain homogenates as substrates. None of the BSE-C samples show *in vitro* propagation. However, 9 of 11 sheep-BSE samples propagated PrP^res^ efficiently. Samples were digested with 200 µg/ml of PK and analyzed by Western blot with monoclonal antibodies D18 (1:5,000). All unseeded samples remained negative. P.: Pooled sample. Dog br.: Undigested *TgDog D163N* brain homogenate. Mw: Molecular weight.

### Attempting to overcome the barrier: from *TgDog D163N* PrP^res^ to *TgDog E163*

We wanted to determine whether, once misfolded by sheep-BSE, the new adapted dog D163N sheep-BSE would be capable of misfolding WT dog PrP^C^ using as PMCA substrates *TgDog* E163 mouse brain homogenates and two different dog brain homogenates coming from different breeds (English Cocker Spaniel and German Wirehaired Pointer). Five rounds of serial PMCA were performed and, even though there was only a single amino acid difference, neither *TgDog* E163 brain homogenate nor dog brain homogenates were able to propagate the adapted dog D163N sheep-BSE, further confirming the reluctance of dog PrP^C^ to misfold and the critical role of the amino acid at position 163 (Fig. 3A). In order to demonstrate that *TgDog D163N*-adapted sheep-BSE retains its propagation capacity and over its original host PrP^C^, cattle brain homogenate was subjected to 5 serial rounds of PMCA, resulting in PrP^res^ with the conserved predominantly diglycosylated band characteristic of this prion strain (Fig. 3B).

**Fig. 3:**
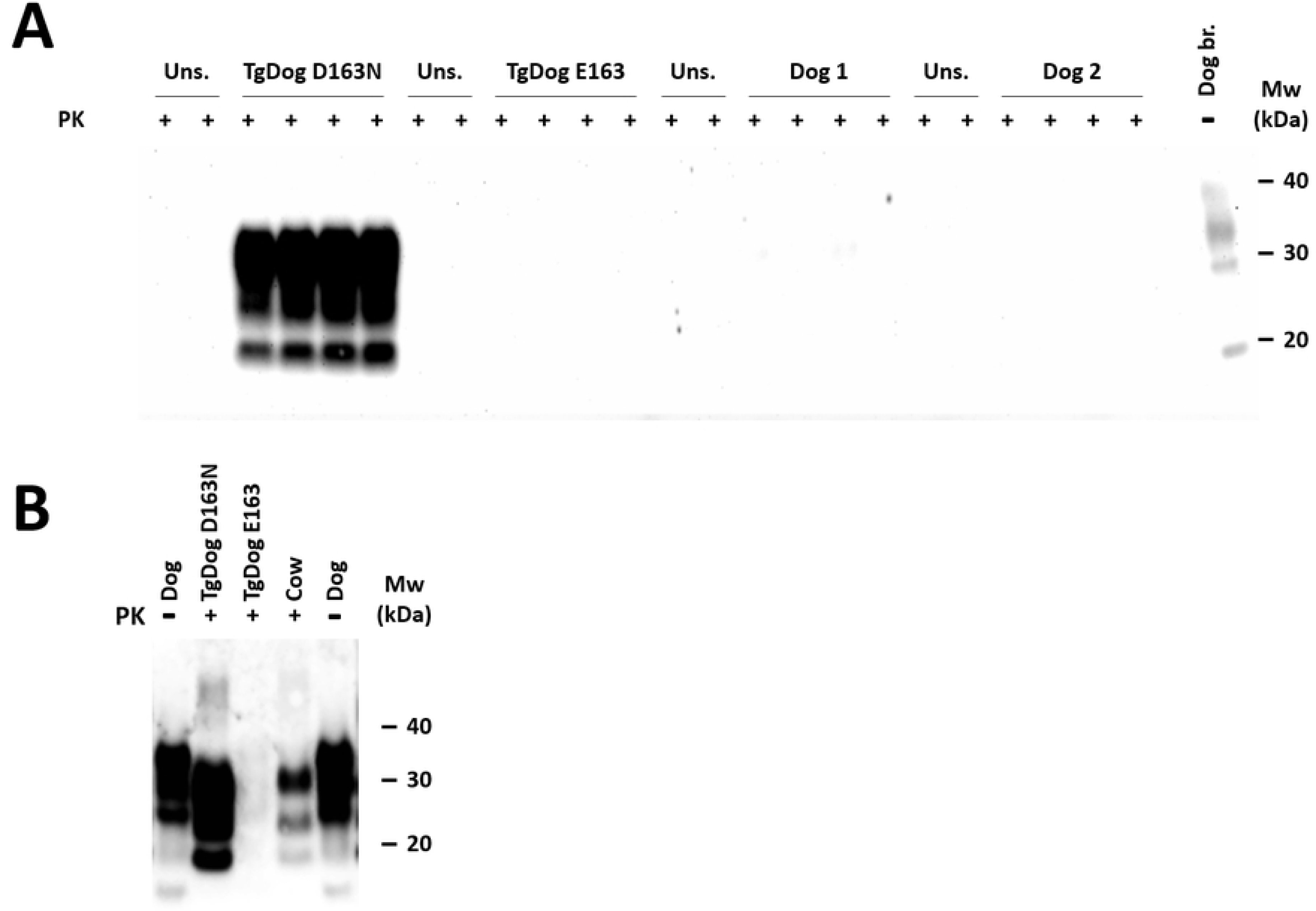
*In vitro* propagation of sheep-BSE inoculated *TgDog D163N* samples using brain-based PMCA. **A.** 3 rounds of serial PMCA using *TgDog D163N*, *TgDog E163* and two different dog breed (English Cocker Spaniel and German Wirehaired Pointer) brain homogenates as substrates. A pool of brain samples from *TgDog D163N* inoculated with sheep-BSE was used as seed in replicates of four through 3 rounds of serial PMCA. Samples were considered positive if a classical PrP^res^ pattern was observed on Western blot. While *TgDog D163N*-based substrate efficiently propagated *in vitro*, neither TgDog E163 nor either of the dog brains used as substrates propagated the *TgDog D163N*-adapted sheep-BSE sample. **B.** In order to demonstrate that the previous (Fig. 3A) *TgDog D163N*-adapted sheep-BSE propagates over a substrate of cow brain (indicating conservation of propagation capacity), this sample was subjected to 5 rounds of serial PMCA using *TgDog D163N*, *TgDog E163* and cow brain homogenates as substrates. Cow brain-based substrate efficiently propagated the *TgDog D163N*-adapted sheep-BSE with a similar pattern characteristic for BSE (predominance of the diglycosylated band) while *TgDog E163* confirmed its inability to propagate this prion strain. Samples were digested with 200 µg/ml of PK and analyzed by Western blot with monoclonal antibody D18 (1:5,000). All unseeded (Uns.) samples remained negative. Dog: Undigested dog (English Cocker Spaniel) brain homogenate. Mw: Molecular weight.

## DISCUSSION

Considerable amounts of data have been generated suggesting that canine PrP^C^ is highly resistant to conformation change to PrP^res^ compared to prion susceptible species[22,32,36,37,41,42]. Our group has now established, both *in vivo* and *in vitro*, that the amino acid residue in position 163 is the key determinant [32,33] and that an aspartic or a glutamic acid in this position (or equivalent in other species) is what conferred resistance to prion infection in the models in which those proteins were expressed[32,34]. Furthermore, these PrP^C^ that are highly resistant to conformation change showed a dominant negative effect when co-expressed with wild type mouse prion protein[32]. Prior to the present report, evidence confirming that murine models expressing wild type canine PrP^C^ were resistant to infection by a panel of prion isolates from different species was lacking and we have addressed this by both bioassays and *in vitro* propagation experiments.

Our conclusions rely on the absence of clinical disease in a single passage in *TgDog E163*. However, when a transmission/species barrier is present, minute amounts of PrP^res^ can be formed on first passage in the absence of any clinical disease or neuropathological lesions at the end of the lifespan of the mouse model. To investigate this phenomenon all challenged animals were not euthanized until the end of their lifespan (or in some cases due to, mostly, age-related intercurrent disease). To ensure the detection of minimal amounts of PrP^res^, pools of inoculated *TgDog* mice brains were subjected to serial PMCA rounds using *TgDog* brain homogenates as a substrate. This technique has been demonstrated to propagate minute amounts of PrP^res^ in the absence of transmission barrier[40]. In all instances no PrP^res^ was present (Supplementary information, Fig. S3) confirming the results obtained in the bioassay.

The total absence of prion infection or *in vitro* propagation with any of the prions used to challenge *TgDog* could be attributed to reasons other than the extreme resistance of canine PrP^C^ to misfolding such as inherent issues with the generation of the transgenic models that may prevent infection. However, this was ruled out through a thorough Western blot analysis of the PrP^C^ of these transgenic animals that showed a virtually identical migration and glycosylation pattern to wild type dog PrP^C^ (albeit expressed at twice the levels) indicating correct posttranslational modifications (Supplementary information, Fig. S1). Additionally, the immunohistochemical localization indicated the correct anatomical expression of the protein on the neuronal cell membrane (Supplementary information, Fig. S1B). Furthermore, the same transgenic generation methodology for obtaining models for other prion disorders has been proven successful previously[28,43]. Altogether, the results support the conclusion that dog PrP^C^ amino acid residue sequence is solely responsible for the complete resistance to prion infection observed in this transgenic model.

The only report unequivocally demonstrating that dog PrP^C^ can be misfolded was achieved only *in vitro*, under highly favorable conditions, by a single prion strain and using dog brain homogenate expressing the 163D polymorphism[22]. However, this *TgDog* model was made prior to understanding the importance of the amino acid residue in position 163. Therefore, in the current work 163E was chosen, as this is exclusive to domestic dogs although both D and E can occur in this position. The 163D polymorphism is present in other canidae species and also in some members of the closely related mustelidae family (wolverine and pine marten, members of the martinae subfamily). Differences in behavior between 163E and 163D containing dog PrP^C^ were explored by cell-based *in vitro* studies and *in silico* analysis of the area containing residue 163 and these revealed differences in the side chain lengths of each residue, the effects of which are unknown[32]. Since the mechanism of action that makes dog PrP^C^ particularly resistant to misfolding might be related to the specific surface charge distribution conferred by these negatively charged amino acids and/or steric hindrance, it is not surprising that the slight differences between the side chains of E and D resulted in small differences in the misfolding capacity of each PrP^C^. The dog prion from the aforementioned study, formed *in vitro* by seeding with BSE (BSE-Dog D163 PrP^res^), was unable to propagate when inoculated intracerebrally in our *TgDog* model indicating that E163 poses a greater limitation for misfolding than D163. However, the BSE-Dog D163 PrP^res^ isolate was propagated efficiently in a TgBov model (BoTg110)[22] suggesting prion amplification without adaptation to the new host but conserving its pathobiological features towards a host with the original cattle PrP^C^, similar to that described previously for other isolate-host combination such as scrapie in TgHorse mice[21].

The possible relevance of the negatively charged amino acid residue in position 163 of dog PrP^C^ to their resistance to prion infection was initially suspected from sequence alignment studies of the *PRNP* of several members of the canidae family with those of susceptible species. These studies revealed that feline PrP was the most similar in terms of amino acid sequence and as domestic cats are susceptible to at least three known prion strains (BSE, CWD and CJD)[26,44–46] the six amino acid difference between canine and feline PrP was studied in detail. The E/D polymorphism in position 163 was highlighted due to its almost exclusive presence in the canidae family and chosen as the most likely candidate for canine resistance to prion disease [32]. Our results with *TgDog*, together with previous reports on canine resistance to prion infection [32–34] clearly identify residue 163 as the strongest effector of the resistance of canine PrP^C^ to misfolding. Therefore, to definitively demonstrate the importance of E or D in position 163, we substituted into the dog PrP the most conserved residue in susceptible species, asparagine, to determine if susceptibility of canine protein to misfolding could be induced. In an experiment conceptually opposite to the one previously conducted with mouse PrP^C^, which was rendered resistant to prion infection by substituting asparagine for aspartic acid at the equivalent position 158[32], the D163N substitution was performed in dog PrP. This eliminates the negative charge and/or steric hindrance that aspartic acid might confer on that region of dog PrP^C^. The resultant transgenic mice could be infected with sheep-BSE inocula. Additionally, *in vitro* amplification using brain homogenates of this *TgDog D163N* model as substrate allowed misfolding of D163N canine PrP^C^ using either sheep-BSE or cattle BSE as seeds. This result strongly supports that amino acid 163 in dog PrP^C^ is the main determinant of its resistance to misfolding by prions. However, since susceptibility was not recovered to all the challenged prion isolates it cannot be regarded as the sole determinant. The prion transmission barrier is assumed to be determined by various factors, including the host PrP sequence and the structure (strain) of the prion. A particular strain will also display a different behavior depending on its own PrP sequence. A clear example of this is BSE, which is more virulent in a sheep PrP^C^ environment than in bovine PrP^C^[47]. Also, the more similar the PrP^C^ sequence is between the host and the inoculated isolate, the lower the transmission barrier should be.

In this regard, it is surprising that the cat-adapted CWD isolate, despite having an amino acid sequence with only a five residues difference from dog D163N PrP^C^ (residues 99, 107, 116, 180 and 188 in dog PrP numbering, some of them identical to the amino acids found in other susceptible species), could not be propagated in *TgDog D163N* substrate which suggest a relevant role of these amino acids in the resistance of this species. It also cannot be excluded that a 2x overexpression of PrP^C^ is not high enough to overcome the transmission barrier.

Interestingly, even sheep-BSE misfolded *TgDog D163N* could not misfold wild type dog PrP^C^ (using *TgDog 163E* brain homogenate as a substrate) in our *in vitro* experiments despite only a single amino acid difference. Again, this suggests a critical role of negatively charged residues at position 163 on PrP misfolding.

In summary, this study provides further experimental evidence that canids, particularly domestic dogs, are the most prion resistant species studied to date and that position 163 in dog PrP^C^ is key in conferring resistance to misfolding thereby establishing this amino acid position coupled with negatively charged residues as a clear therapeutic target for prion diseases.

## Materials and methods

### Preparation of inocula for prion propagation studies

Brain homogenates (10^−1^ in PBS) for use as seeds for PMCA or direct intracerebral inoculation were prepared manually using a glazed mortar and pestle from brains of animals clinically affected by various TSE: BSE-C and scrapie (SSBP/1) were supplied by Animal & Plant Heath Agency (UK), BSE-L field cases were supplied by Centro di Referenza Nazionale per le Encefalopatie Animali (Turin, Italy), CWD from the thalamus area of the brain of a female mule deer, genotype 225SS, infected with CWD (04–22412WSV2 EJW/JEJ), supplied by Department of Veterinary Sciences (University of Wyoming, Laramie, WY, USA), feline CWD from an experimental case of CWD infection in a domestic cat was supplied by Department of Microbiology, Immunology and Pathology, Colorado State University (Fort Collins, Colorado, USA)[48], and Sheep-BSE was supplied by Ecole Nationale Vétérinaire (Toulouse, France). The atypical scrapie isolate was obtained from a field case diagnosed in the PRIOCAT laboratory, CReSA-IRTA (Barcelona, Spain). BSE DoD163 PrP^res^ was generated previously by PMCA using cattle BSE as the seed[22].

### Generation of *in vitro* PrP^res^ by serial PMCA

The *in vitro* prion replication and PrP^res^ detection of amplified samples was performed as described previously with minor modifications[49]. Briefly, brains used for substrate were perfused using PBS + 5 mM EDTA and the blood-depleted brains were frozen immediately until required for preparing the 10 % brain homogenates (PBS + NaCl 0.15 M + 1% Triton X-100). Brain homogenates (50–60 μl of 10 %), either unseeded or seeded with the corresponding prion isolate/strain, were loaded into 0.2-ml PCR tubes and placed into a sonicating water bath at 37–38 °C without shaking. Tubes were positioned on an adaptor placed on the plate holder of the sonicator (model S-700MPX, QSonica, Newtown, CT, USA) and subjected to incubation cycles of 30 min followed by a 20 s pulse of 150–220 watts sonication at 70–90 % amplitude. Serial rounds of PMCA consisted of 24-48h of standard PMCA followed by serial *in vitro* 1:10 passages in fresh 10 % brain homogenate substrate. An equivalent number of unseeded (4 duplicates) tubes containing the corresponding brain substrate were subjected to the same number of rounds of PMCA in order to monitor for cross-contamination and/or the generation of spontaneous PrP^res^.

### Biochemical characterization of *in vitro*- and *in vivo*-generated prion strains

PMCA treated samples were incubated with 85–200 μg/ml of protease K (PK) for 1 h at 42 °C with shaking (450 rpm) as described previously[50]. Digestion was stopped by adding electrophoresis Laemmli loading buffer and the samples were analyzed by Western blotting.

### Generation of *TgDog E163* and *TgDog D163N* mice

After isolation by PCR amplification from genomic DNA extracted using GeneJET™ Genomic DNA Purification Kit (Fermentas) from a E163 dog tissue sample using 5’ GGGGGAATTCATCATGGTGAAAAGCCACATAGGCG 3’ and 5’ GGGCGGGCGGCCGCTCATCCCACTATCAAGAGAATG 3’ as primers, the open reading frame (ORF) of the E163 dog *PRNP* gene was cloned into the pGEM-T vector (Promega). In the same way, the ORF of D163 dog *PRNP* was isolated from the genomic DNA extracted from the tissue sample of a dog bearing D163 polymorphism using the same primers and cloned into pGEM-T vector. The dog E163-PrP ORF was excised from the cloning vector by using the restriction enzymes BsiWI (Thermo Fisher Scientific Inc.) and FseI (New England Biolabs Ltd.) and then inserted into a modified version of MoPrP.Xho vector[51] as described previously[38], which was also digested with BstWI and FseI. This vector contains the murine PrP promoter and exon-1, intron-1, exon-2 and 3’ untranslated sequences. The genetic construct containing the dog D163N substitution was carried out by two-step PCR site-directed mutagenesis using pGEM dog D163 as template, using primers 5’ GAACATGTACCGCTACCCCAACCAAGTATACTACCGG 3’ with 5’ GGGCGGGCGGCCGCTCATCCCACTATCAAGAGAATG 3’ and 5’ CCGGTAGTATACTTGGTTGGGGTAGCGGTACATGTTC 3’ with 5’ GGGGGAATTCATCATGGTGAAAAGCCACATAGGCG 3’. Then using the previous fragments as templates and primers 5’ GGGGGAATTCATCATGGTGAAAAGCCACATAGGCG 3’ and 5’ GGGCGGGCGGCCGCTCATCCCACTATCAAGAGAATG 3’, the dog D163N-PrP ORF was generated and cloned into the pGEM-T vector (Promega). It was also excised from the cloning vector using restriction enzymes BsiWI (Thermo Fisher Scientific Inc.) and FseI (New England Biolabs Ltd.), and then inserted into a modified version of MoPrP.Xho vector. Both transgenes were excised using NotI and purified with an Invisorb Spin DNA Extraction Kit (Inviteck) according to the manufacturer recommendations.

Transgenic mouse founders were generated by microinjection of DNA into pronuclei following standard procedures[29]. DNA extracted from tail biopsies was analyzed by PCR using specific primers for the mouse exon 2 and 3’ untranslated sequences (5’ GAACTGAACCATTTCAACCGAG 3’ and 5’ AGAGCTACAGGTGGATAACC 3’). Those which tested positive were bred to mice null for the mouse *PRNP* gene in order to avoid endogenous expression of mouse prion protein. Absence of the mouse endogenous *PRNP* was assessed using the following primers: 5’ ATGGCGAACCTTGGCTACTGGC 3’ and 5’ GATTATGGGTACCCCCTCCTTGG 3’. The dog PrP expression levels of brain homogenates from transgenic mouse founders were determined by Western blot using anti-PrP mAb D18 [52] and compared with the PrP expression levels from different dog brain homogenates.

The international code to identify these transgenic mouse lines are STOCK-Prnptm2Edin Tg(moPrpn dogPrP)14Bps and 129OLA-Prnptm2Edin-Tg(mPrpn-dogPrPD163N)1Sala although throughout the paper they are referred to as *TgDog E163* and *TgDog D163N* mice, respectively.

### *TgDog E163* and *TgDog D163N* mice inoculation

Mice of 42-56 days of age were intracerebrally inoculated under gaseous anesthesia (Isoflurane) through the right parietal bone. A 50 µl SGC precision syringe was used with a 25 G gauge needle and coupled to a repeatability adaptor fixed at 20 µl. A dose of buprenorphine was subcutaneously injected before recovery to consciousness to reduce post-inoculation pain. Mice were kept in a controlled environment at a room temperature of 22 °C, 12 h light-darkness cycle and 60 % relative humidity in HEPA filtered cages (both air inflow and extraction) in ventilated racks. The mice were fed *ad libitum*, observed daily and their clinical status assessed twice a week. The presence of ten different TSE-associated clinical signs [53] was scored. Positive TSE diagnosis relied principally on the detection of PrP^res^ (either by immunohistochemistry and/or western blotting or ELISA) and associated spongiform changes on stained histological sections (see below) of the brain parenchyma.

### Sample processing and general procedures

When the clinical end-point criteria were reached mice were euthanized by decapitation. The brain was extracted immediately and placed into 10 % phosphate buffered formalin. Transversal sections of the brain were performed at the levels of the medulla oblongata, piriform cortex and optic chiasm. Samples were embedded in paraffin-wax after dehydration through increasing alcohol concentrations and xylene. Four micrometer sections were mounted on glass microscope slides and stained with hematoxylin and eosin for morphological evaluation. Additional sections were mounted in 3-trietoxysilil-propilamine-coated glass microscope slides for immunohistochemistry. The spinal cord and a partial section of the frontal cortex, including the olfactory bulbs, were separated prior to fixation and kept frozen for biochemical analysis.

### Immunohistochemistry

Immunohistochemistry (IHC) for detection of PrP^res^ was performed as described previously [54]. Briefly, deparaffinized sections were subjected to epitope unmasking treatments: immersed in formic acid and boiled at low pH (6.15) in a pressure cooker and pre-treated with proteinase K. Endogenous peroxidases were blocked by immersion in a 3 % H_2_O_2_ in methanol solution. Sections were then incubated overnight with anti-PrP MAb 6H4 primary antibody (1:2,000, Prionics AG) and subsequently visualized using the DAKO Goat anti-mouse EnVision system (Ref. K400111/0) and 3,3’-diaminobenzidine as the chromogen substrate. As a background control, incubation with the primary antibody was omitted.

## Ethics Statement

All experiments involving *TgDog* animals were approved by the animal experimentation ethics committee of the Autonomous University of Barcelona (Reference number: 585-3487) in agreement with Article 28, sections a), b), c) and d) of the “Real Decreto 214/1997 de 30 de Julio” and the European Directive 86/609/CEE and the European Council Guidelines included in the European Convention for the Protection of Vertebrate Animals used for Experimental and Other Scientific Purposes.

All experiments involving *TgDog D163N* animals were approved by the Ethical Committee on Animal Welfare of the Laboratorio Central de Veterinaria (project code assigned by the Ethical Committee CEBA-07/2010) and also in agreement with the aforementioned European legislation and the Spanish Legislative Decree “Real Decreto 1201/2005 de 10 de Octubre”.

## Acknowledgments

The authors would like to thank the following for their support: IKERBasque foundation, vivarium and maintenance from CIC bioGUNE and Patricia Piñeiro for technical support. Sierra Espinar, Marta Valle and Mariano Moreno for technical support, and IRTA-CReSA’s biocontainment unit staff for care and maintenance of the animals. Olivier Andréoletti, Jean Jewell and Fabrizio Tagliavini for the sheep-BSE, CWD and BSE-L brain tissue samples, respectively. Glenn Telling and Jifeng Bian for the plasmid phggPrP-MCS. Dr. Mark P. Dagleish (Moredun Research Institute) for useful discussion and advice.

## Supporting information legends

**Fig. S1: PrP expression levels in *TgDog E163* animals compared to PrP expression levels of normal dog brain by Western blot**.

**A:** 10% brain homogenates from *TgDog E163* mouse and dog were diluted 1:20, 1:40, 1:80, 1:160, 1:320 and 1:640 and analyzed by Western blot using monoclonal antibody D18 (1:5,000). The PrP expression levels of *TgDog E163* were approximately double compared to PrP^C^ levels in dog brain, based on signal intensity. Notice that the glycosylation pattern is maintained between the transgenic mice and the dog indicating correct posttranslational processing of the PrP^C^ in the mouse. Mw: Molecular weight. **B**: **Immunohistochemical analysis of PrP^C^ expression in *TgDog E163* and *TgDog D163N* compared to C57BL/6 mice.** Cerebral cortex sections from *TgDog E163* and C57BL/6 mice were used to compare the localization of PrP^C^ expression. A diffuse neuropil immunolabeling (corresponding to PrP^C^ on the dendrite cell membrane) and absence of labeling within the pericarion were observed. PrP^C^ immunolabeling from *TgDog E163* brain was comparable to that found in WT (C57BL/6) brains, revealing a normal synaptic staining. Nuclear staining is artefactual. Samples were immunostained using 6C2 (1:100) monoclonal antibody. Bar: 25 μm.

**Fig. S2: *In vitro* propagation of different prion isolates using *TgDog E163* brain-based PMCA.**

Rounds (R1-R10) of serial PMCA using *TgDog E163* (line 014) transgenic mouse brain homogenates as substrates. The prion isolates: BSE-C, sheep-BSE, Scrapie (SSBP/1), BSE-L, atypical scrapie, mule deer CWD, cat CWD, BSE-dog D163 (*in vitro* generated) or unseeded, were used as seeds in replicates of four through 10 rounds of serial PMCA. Samples were considered positive if a classical PrP^res^ pattern was observed on Western blot. None of the isolates propagated over the *TgDog E163* substrate.

**Fig. S3: *In vitro* propagation of different prion-infected samples using *TgDog E163* brain-based PMCA.**

Rounds (R1-R6) of serial PMCA using *TgDog E163* (line 014) transgenic mouse brain homogenates as substrates. The brains from *TgDog E163* mice inoculated with the different prion isolates (see supplementary figure 2) were pooled and were used as seeds in replicates of four through 6 rounds of serial PMCA. Samples were considered positive if a classical PrP^res^ pattern was observed on Western blot. None of the samples propagated over the *TgDog E163* substrate failing to demonstrate propagation of even minute amounts of misfolded prion protein.

**Fig. S4: PrP expression levels in *TgDog D163N* animals compared to PrP expression levels of dog by Western blot.**

**A:** 10% brain homogenates from *TgDog D163N* mouse and dog were diluted 1:20, 1:40, 1:80, 1:160, 1:320 and 1:640 and analyzed by Western blot using monoclonal antibody D18 (1:5,000). The PrP expression levels of *TgDog D163N* were approximately double to PrP^C^ levels in dog brain based on signal intensity. Notice that the glycosylation pattern was maintained between the transgenic mice and the dog indicating correct posttranslational processing of the PrP^C^ in the mouse. Mw: Molecular weight. **B**: **Immunohistochemical analysis of PrP^C^ expression in *TgDog E163* and *TgDog D163N* compared to C57BL/6 mice.** Cerebral cortex sections from *TgDog E163* and *TgDog D163N* and C57BL/6 mice were used to compare the localization of PrP^C^ expression. A fine granular neuropil immunolabeling (corresponding to PrP^C^ on the dendrite cell membrane) and absence of labeling within the pericarion were observed. PrP^C^ immunolabeling from *TgDog D163N* brains was comparable to that found in WT (C57BL/6) brains, revealing a normal synaptic staining. Nuclear staining is artefactual. Samples were immunostained using 6C2 (1:100) monoclonal antibody. Bar: 25 μm.

**Fig. S5: *In vitro* propagation of different prion isolates using *TgDog D163N* brain-based PMCA.**

Rounds (R1-R10) of serial PMCA using *TgDog D163N* (line 483) transgenic mouse brain homogenates as substrates. The prion isolates: BSE-C, sheep-BSE, Scrapie (SSBP/1), BSE-L, atypical scrapie, mule deer CWD, cat CWD or unseeded, were used as seeds in replicates of four through 10 rounds of serial PMCA. Samples were considered positive if a classical PrP^res^ pattern was observed on Western blot. Cattle BSE-C and sheep-BSE propagated over the *TgDog D163N* substrate but none of the other prion isolates resulted in any propagation.

**Fig. S6: *In vitro* propagation of different prion isolates using *TgDog D163N* brain-based PMCA.**

Rounds (R1-R6) of serial PMCA using *TgDog D163N* (line 483) transgenic mouse brain homogenates as substrates. The brains from *TgDog D163* mice inoculated with the different prion isolates (see supplementary figure 5) were pooled and used as seeds in replicates of four through 6 rounds of serial PMCA. Samples were considered positive if a classical PrP^res^ pattern was observed on Western blot. Just sheep-BSE propagated over the *TgDog D163N* substrate.

